# Feedbacks between size and density determine rapid eco-phenotypic dynamics

**DOI:** 10.1101/2021.07.15.452551

**Authors:** Jean Philippe Gibert, Ze-Yi Han, Daniel J Wieczynski, Samantha Votzke, Andrea Yammine

**Affiliations:** Duke University

**Keywords:** Plasticity, Eco-evolutionary Dynamics, Traits, Phenotypes, Phenotypic change

## Abstract

1. Body size is a fundamental trait linked to many ecological processes—from individuals to ecosystems. Although the effects of body size on metabolism are well-known, the potential reciprocal effects of body size and density are less clear. Specifically, 1) whether changes in body size or density more strongly influence the other and 2) whether coupled rapid changes in body size and density are due to plasticity, rapid evolutionary change, or a combination of both.
2. Here, we address these two issues by experimentally tracking population density and mean body size in the protist *Tetrahymena pyriformis* as it grows from low density to carrying capacity. We then use *Convergent Cross Mapping* time series analyses to infer the direction, magnitude, and causality of the link between body size and ecological dynamics. We confirm the results of our analysis by experimentally manipulating body size and density while keeping the other constant. Last, we fit mathematical models to our experimental time series that account for purely plastic change in body size, rapid evolution in size, or a combination of both, to gain insight into the processes that most likely explain the observed dynamics.
3. Our results indicate that changes in body size more strongly influence changes in density than the other way around, but also show that there is reciprocity in this effect (i.e., a *feedback*). We show that a model that only accounts for purely plastic change in size most parsimoniously explains observed, coupled phenotypic and ecological dynamics.
4. Together, these results suggest 1) that body size can shift dramatically through plasticity, well within ecological timescales, 2) that rapid changes in body size may have a larger effect on ecological dynamics than the reverse, but 3) phenotypic and ecological dynamics influence each as populations grow. Overall, we show that rapid plastic changes in functional traits like body size may play a fundamental –but currently unrecognized– role in familiar ecological processes such as logistic population growth.

## INTRODUCTION

Body size influences organismal energetic demands (Gillooly *et al*. 2001; Brown *et al*. 2004), diet breadth (Wasserman & Mitter 1978; Gravel *et al*. 2013), the strength of ecological interactions (Berlow *et al*. 2009), trophic cascades (DeLong *et al*. 2015), and food web structure (Brose *et al*. 2006; Riede *et al*. 2011; Gravel *et al*. 2013; Gibert & DeLong 2014), all of which have ecosystem-level consequences (Anderson-Teixeira, Vitousek & Brown 2008; Trebilco *et al*. 2013). Because of its myriad ecological consequences—from individuals (e.g., (Glasheen & McMahon 1996; Hurlbert, Ballantyne IV & Powell 2008; Pavković-Lučić & Kekić 2013)) to ecosystems (e.g., (Brown *et al*. 2004; Riede *et al*. 2011; Trebilco *et al*. 2013))—body size is one of the most important functional traits.

Body size can fuel or limit population growth through well-known relationships with demographic parameters like carrying capacity (K) and the intrinsic growth rate (r) (Damuth 1981; Savage *et al*. 2004; DeLong *et al*. 2015). For example, smaller organisms typically reproduce faster and can reach higher carrying capacities than larger ones, but also have higher mortality rates, which leads to faster population turnover (e.g., (Huryn & Benke 2007)). As a consequence, body size often determines how populations grow, and hence, ecological dynamics. On the flip side, ecological dynamics can themselves influence body size, although these effects are not well understood. For example, individual size is under physiological regulatory control (Davidowitz, Roff & Nijhout 2005; Chelini, Delong & Hebets 2019) and therefore responds to changes in resource levels (Marañón *et al*. 2013). While resource levels can vary independently of population density (Holt 2008; Fey, Gibert & Siepielski 2019; Nguyen, Lara-Gutiérrez & Stocker 2020), high density leads to crowding and increasing intra-specific competition (Gavina *et al*. 2018). Stronger competition in turn reduces resource availability at the individual level, which may result in stunted growth and smaller body sizes (e.g., (Vanni *et al*. 2009)). Consequently, while body size can, and often does, influence ecological dynamics, ecological dynamics can also influence body size. Which one more strongly influences the other, however, is not well understood, among other reasons because both population density and body size can change dramatically over time (DeLong, Hanley & Vasseur 2014; Clements & Ozgul 2016).

Due to either natural or sexual selection (Preziosi & Fairbairn 2000; Chelini, Delong & Hebets 2019), body size can change evolutionarily (Hairston *et al*. 2005; DeLong *et al*. 2016). For example, predation selects for smaller but faster-growing prey in pitcher plant inquiline communities (terHorst, Miller & Levitan 2010) and for shorter developmental times that result in smaller individuals in mayflies (Peckarsky *et al*. 2001). Selection can act on traits rather quickly (Thompson 1998; Hairston *et al*. 2005) and rapid evolutionary change has been shown to influence ecological dynamics as they unfold (e.g., (Becks *et al*. 2012; Rudman *et al*. 2018; Schaffner *et al*. 2019)), while ecological dynamics, in turn, influence the pace and direction of evolutionary change (e.g., (Cortez 2016; DeLong & Gibert 2016; Frickel, Sieber & Becks 2016; Gibert & Yeakel 2019)). Therefore, selection imposed by ecological dynamics may influence body size, whose rapid change can alter ecological dynamics. Because studying rapid evolutionary change in body size is unfeasible for most organisms, it is unclear how pervasive these processes are in nature.

In addition to evolution, body size can change within generations through plasticity (David, Legout & Moreteau 2006; Ghosh, Testa & Shingleton 2013; Lafuente, Duneau & Beldade 2018; Cameron *et al*. 2020). For example, organisms grow in size throughout ontogeny and the environment often influences those ontogenetic trajectories, leading to plastic variation in body size (Lafuente, Duneau & Beldade 2018; Chelini, Delong & Hebets 2019). Epigenetic DNA modifications can also result in rapid phenotypic change from one generation to the next in response to shifts in biotic or abiotic conditions (e.g., maternal effects, (Roach & Wulff 1987; Galloway & Etterson 2007)). Teasing apart the contributions of plastic and evolutionary processes on body size has been the subject of great scientific interest (Amarillo-Suarez, Stillwell & Fox 2011; Walczyńska, Franch-Gras & Serra 2017; Lafuente, Duneau & Beldade 2018; Yengo *et al*. 2018; Cox *et al*. 2019). However, whether plasticity or evolution more strongly influences rapid changes in body size, especially when this is coupled to rapidly shifting ecological dynamics, is not well understood (e.g., (DeLong, Hanley & Vasseur 2014)). Among other reasons, this is because teasing apart plastic and evolutionary change over short periods of time is challenging, even when sufficiently long time series are available (Ellner, Geber & Hairston 2011). As body size mediates ecological interactions and processes, and can change on ecological timescales, it is important to understand, track, and predict such change.

Here, we address these gaps by answering the following questions: 1) Do rapid changes in body size more strongly influence population dynamics (i.e., changes in density over time), or do population dynamics more strongly influence changes in body size? and 2) Are observed changes in body size most consistent with a model of plasticity, rapid evolution, or one that accounts for both plastic and evolutionary change? To address the first question, we track changes in the density and average body size of multiple experimental populations of the protist *Tetrahymena pyriformis*, then use time series analysis to infer causality. In protists, we expect changes in body size to be at least partly caused by plasticity because reproduction (cell division) is tightly linked to ontogenetic changes in body size (cells grow then divide when a critical size is attained). However, *T. pyriformis* also reproduces extremely fast (∼4 generations per day) and exhibits wide standing variation in body size (Wieczynski *et al*. 2021), so rapid evolutionary change is also possible. Therefore, to distinguish the impacts of plasticity and evolution on changes in body size, we fit alternative mathematical models (Plasticity Model, Eco-Evolutionary Model, and Plasticity + Eco-Evo Model) to our experimental time series and use model selection to infer which one best explains our data. Our results show that rapid, purely plastic changes in body size more strongly influence changes in density than the other way around, thus suggesting that plastic phenotypic change may be integral to ecological dynamics.

## METHODS

### Microcosm growth assays

We tracked changes in abundance and body size in the protist *Tetrahymena pyriformis* for 14 days. To do so, we set up 6 experimental microcosms in 250 mL autoclaved borosilicate jars filled with 200 mL of Carolina protist pellet media (1L of autoclaved DI water per pellet) inoculated with pond bacteria from Duke Forest (Gate 9/Wilbur pond, Lat=36.02°, Long=-78.99°, Durham, NC) and a wheat seed as a carbon source. Microcosms were initialized at densities of 10 ind/mL and incubated in temperature (22°C) and humidity-controlled (65% humidity) growth chambers (Percival AL-22L2, Percival Scientific, Perry, Iowa) on a 12hr night/day cycle. Densities (ind/mL) and trait dynamics were tracked daily for two weeks through fluid imaging of 1 mL subsamples of each microcosm (Fig 1a, FlowCam, Fluid Imaging Technologies, Scarborough, ME, USA). The FlowCam can image particles ranging from 5-10 μm to 2mm in length. Cell images were automatically sorted and measured by the FlowCam’s proprietary software yielding individual-level data on 150k cells over 14 days, giving our experiment unparalleled insight into how density and body size changed together over the course of this experiment. Using these data, density was quantified as a simple cell count per volume sampled and body size was quantified as the volume of a spheroid, in*μ*m^3^. Last, we quantified changes in total biomass, measured as the sum of the mass of all individual cells in a sample (in grams, *g*, obtained by converting protist volumes estimated by the FlowCam from μm^3^ to cm^3^ and assuming that the density of protists equals that of water, i.e., 1g/cm^3^). Neither water nor nutrients were replaced throughout the course of this experiment.

**Fig 1:**
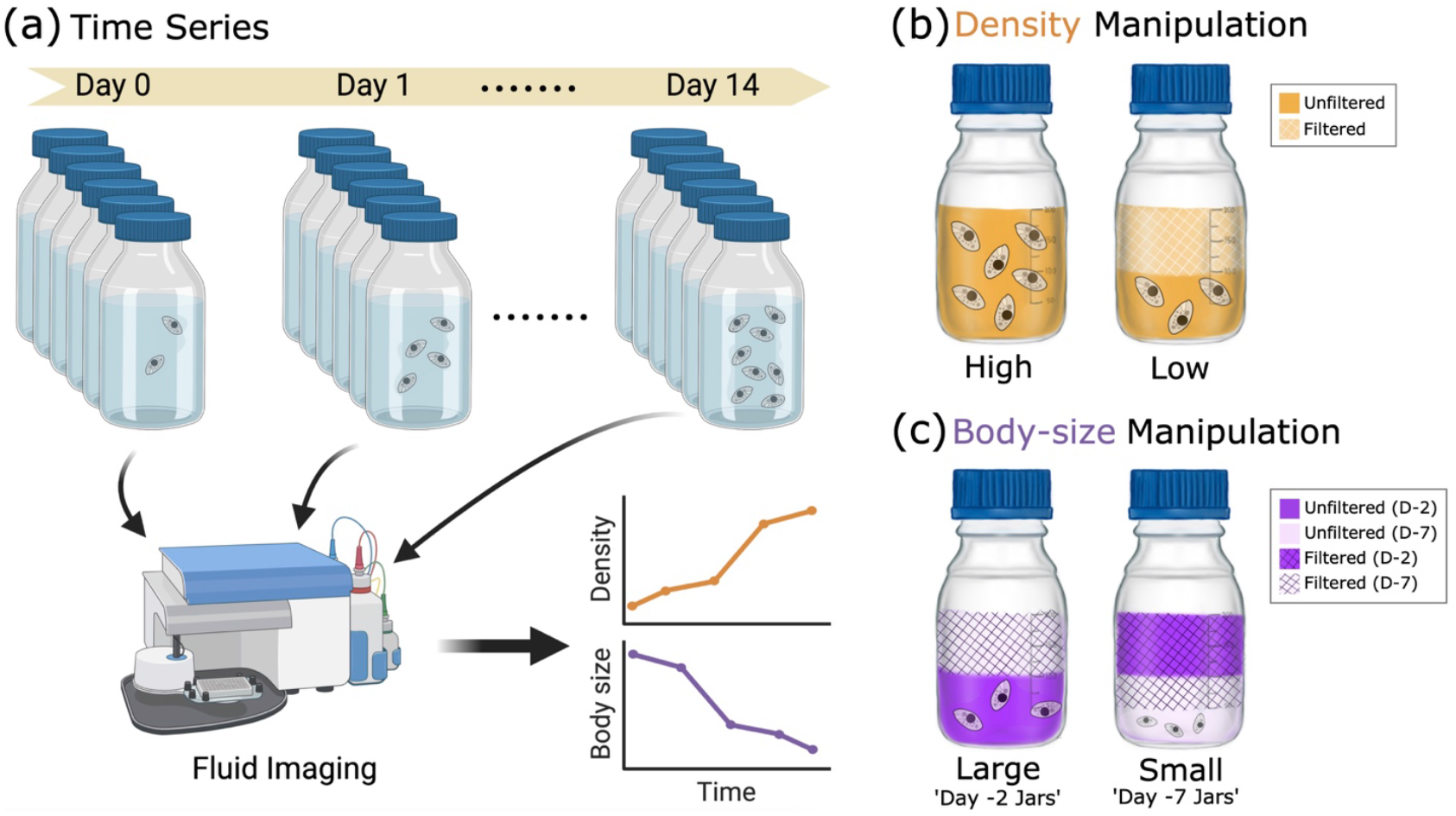
(a) Depiction of experimental procedure: microcosms were initialized at 10ind/mL and each microcosm was replicated six times. Population density and cell size were recorded daily for 14 days thereafter using fluid imaging. (b) Experimental setup to manipulate density while controlling for body size. This involved filtering the Tetrahymena out of 100mL of media in half of the experimental jars. (c) Experimental setup to manipulate body size while controlling for density. This involved the filtration of 100mL of media of Day-2 (D-2) and Day-7 (D-7) jars, which was then added to jars of the other group (ensuring equal resources and media). An extra filtration step ensured that the density in 100mL of Day-2 jars equaled the density in 100mL of Day-7 jars. To find the volume (!) to be filtered from 100mL of unfiltered Day-7 media (then returned to Day-7 jars), we noticed that 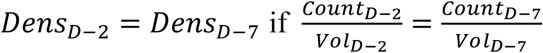, and that *vol*_*D*−2_=100*mL* and *vol*_*D*−7_=100 *mL* − *x*. Solving for *x* yielded how much volume had to be filtered and returned to the same jars. This extra filtering step was done first and the filtrate was set aside before the other two steps.

### Time-series analysis

To assess whether change in body size more strongly influenced changes in density, or vice versa, we used *Convergent Cross Mapping* (or CCM, (Sugihara *et al*. 2012; Rogers *et al*. 2020)) on the density and body size time series. Conceptually, CCM quantifies the degree to which one time series causally influences another one by estimating how much information of the one is contained in the other (Takens 1981; Sugihara *et al*. 2012). If a variable X causally influences another variable Y, but Y does not influence X, Y should contain information about X, but not the other way around. CCM does that by quantifying whether variable X can be predicted from the time series of Y (and vice-versa) for subsets of the time series of increasing length (this procedure is called ‘cross-mapping’). If X more strongly influences changes in Y than the other way around, it also means Y responds to X more strongly than X responds to Y. If the effect of X on Y is causal –as opposed to there being simple correlation with an unobserved variable Z– the ability to predict Y from X should increase with library size, while the error associated with that prediction should decline. If predictability does not change with library size, there is correlation, but not causation (e.g., variable X and Y could be responding to a third unobserved variable Z instead of to each other, leading to spurious correlation between the two, (Sugihara *et al*. 2012)).

To perform the analysis, we used R package multispatialCCM v1.0 (Clark *et al*. 2015), which works on replicated time series. In a nutshell, the procedure operates as follows: first, the algorithm does state-space reconstruction using ‘delay-embedding’ (Sugihara *et al*. 2012). That is, it attempts to reconstruct the manifold of the system (i.e., the collection of all states taken by all variables for all time points) using lagged versions of each variable, one at a time. The number of lagged versions of each time series needed for this reconstruction is called the ‘Embedding Dimension’, *E* (Sugihara & May 1990; Sugihara *et al*. 2012). A value of E much larger than the number of observed variables suggests effects of other non-observed variables on the dynamics of the system (Sugihara *et al*. 2012). Second, the ‘multispatialCCM’ version of the CCM algorithm takes bootstrap pseudo-replicates (n=800) of varying size (i.e., library size, ranging from E to E*n, where n is the number of replicated time series) across replicates (Clark *et al*. 2015). Third, it uses those bootstrapped time series to ‘cross-map’, that is, to predict the values of one state variable based on exponentially weighted values of the reconstructed manifold of the system using the other state variable (i.e., ‘predicting X based on information contained in Y’). By quantifying the correlation coefficient (or the *predictability* coefficient) of the observed and predicted values, CCM produces a measure of how strong of a causal effect one variable has on the other (Sugihara *et al*. 2012), if such an effect exists and is indeed causal.

Multiple previous studies have already shown how well CCM infers causation in different ecological systems and environmental conditions (Sugihara *et al*. 2012; Clark *et al*. 2015; Karakoç, Clark & Chatzinotas 2020; Kondoh *et al*. 2020; Rogers *et al*. 2020).

### Experimental manipulations of size and density

In addition to our time series analysis, we experimentally manipulated density and size, while controlling for the other variable, to assess whether possible effects of size on density and vice-versa could be detected experimentally.

### Experiment 1: effect of density on body size

To manipulate density while keeping body size constant, we started 30 microcosms (as detailed in *Microcosm growth assays* subsection) populated with *T. pyriformis* at an initial density of 10ind/mL, at Day -2. At Day 0, we filtered half of the volume (100 mL) in 15 of those microcosms using Whatman GF/A filters, which have a pore size small enough to filter out the protists, but large enough to allow the bacteria protists feed on to pass through. The original microcosms were then replenished with the filtered water with bacteria (but no protists). This procedure halved the density of 15 out of the initial 30 jars (Fig 1b) while keeping the size distribution of *T. pyriformis*, growth medium, and number of bacteria, the same, in ‘low’ (jars with half filtered, half unfiltered growth medium) and ‘high’ density treatments (jars with unfiltered growth medium).

### Experiment 2: effect of body size on density

To manipulate body size while keeping density constant, we started 15 microcosms (as described before) at Day -7 (‘Day -7 jars’ henceforth), and another 15 microcosms at Day -2 (‘Day -2 jars’ henceforth). At Day 0, we removed and filtered, as before, half of the volume (100 mL) in all jars. We then replenished all jars with filtered medium from jars in the other group, so all jars contained a mixture of equal parts medium and bacteria (resources) from Day -2 and Day -7 jars (Fig 1c). Simultaneously, we filtered an additional amount of medium from Day -7 jars to ensure that the cell density in this group, once all growth medium had been added, matched that of day -2 jars (Fig 1c). This was done by calculating the volume of growth medium of the original Day -7 jars that needed to be filtered based on the observed density (after manipulation) in Day -2 jars. Because *T. pyriformis* decays in size as it grows to carrying capacity (Fig 2a, b), Day -7 jars contained, on average, smaller individuals, Day -2 jars contained relatively larger individuals (Fig 1c), but both groups had the same population density, medium, and bacterial density (resources).

**Fig 2:**
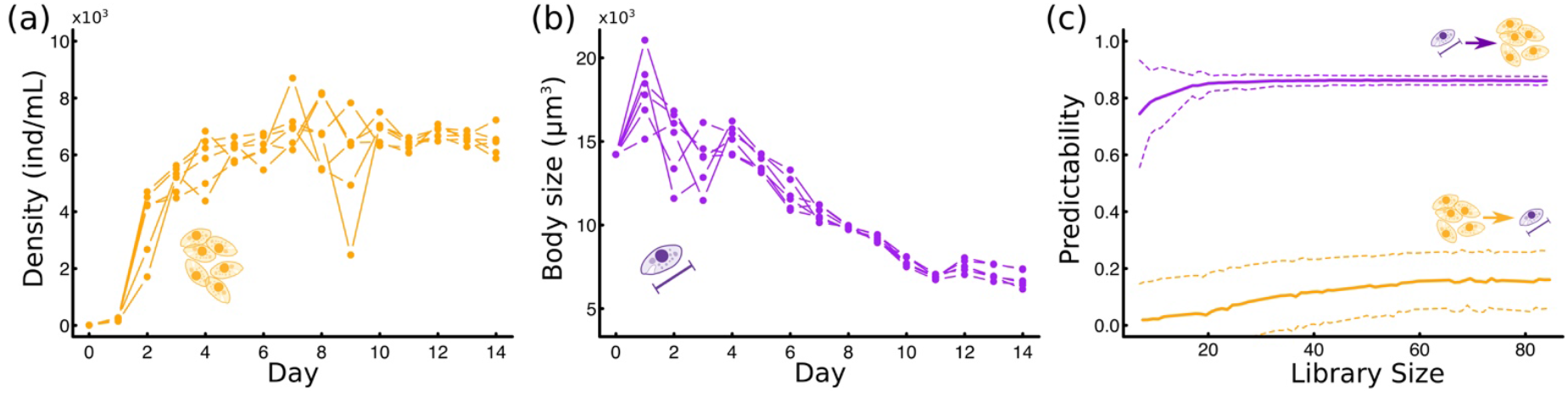
(a) Time series of Tetrahymena density. (b) Time series of mean Tetrahymena body size. (c) Convergent Cross Mapping predictability plot against library size (i.e., length of the time series used for analysis), using body size to predict density (purple) or density to predict body size (yellow), repeated 1000 times. Solid line indicates mean values and dashed lines indicate standard deviation of the mean.

We used the FlowCam to quantify body size and density in Day 0 (the day the manipulations where made) and Day 2 (i.e., two days later). In both experiments, we expected density and size to change over time (from Day 0 to Day 2), meaning that statistically speaking, we expect time to influence both density and size. However, if either size or density influence the other, we also expect the interaction between time and density (experiment 1), or time and size (experiment 2), to be significant. In that case, the interaction term indicates by how much a difference in starting density or size influences the change in the response variable as time elapses. A large (small) interaction term would indicate a large (small) effect of the initial difference in either size or density on the response variable. We tested for these interactions using linear models in R v3.6.1 (R Core Team 2013) with either 1) body size as a response variable and density, day, and their interaction as predictors (experiment 1) or 2) density as a response variable and body size, day, and their interaction as predictors (experiment 2). To compare the effect sizes of density on size, and size on density, we standardized all variables in R package *effectsize* v0.6.01 (Ben-Shachar, Lüdecke & Makowski 2020) by re-centering and re-scaling variables to a normal distribution with mean equal to zero and standard deviation equal to 1 (Gelman 2008).

### Mathematical models

While others have argued that laboratory cultures should have low levels of heritable genetic variation (DeLong, Hanley & Vasseur 2014), *T. pyriformis* has mechanisms to maintain larger than expected levels of genetic variation (Dimond & Zufall 2016). Because of this, and to assess whether observed changes in body size were more likely due to plasticity or rapid evolution, we fitted two possible models that track change in the abundance and average body size of a population, *N*, as it grows logistically towards a carrying capacity *K* with intrinsic growth rate *r*. Following previous work (Abrams 1977; Abreu *et al*. 2019; Lax, Abreu & Gore 2020; Wieczynski *et al*. 2021), we included an additional mortality term in the ecological dynamics, *mN*, to account for regular loss of individuals from the population through sampling. This additional mortality term has been shown to better describe the ecological dynamics of a microbial microcosm with frequent sampling, like ours (Abreu *et al*. 2019; Lax, Abreu & Gore 2020). Furthermore, we assumed that the intrinsic growth rate and the carrying capacity were functions of (average) body size, *M*, of the forms *r*=*aM*^*α*^and *k*=*bM*^*γ*^, following well-known allometric relationships (Damuth 1981; Savage *et al*. 2004). Taken together, the ecological baseline model was thus written as:

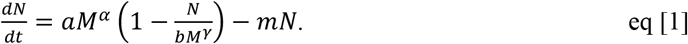

To incorporate coupling between ecological dynamics (changes in N) and both plastic and evolutionary changes in *M*, we used three alternative model formulations to track changes in *M*: the first assumed that only plastic change in *M* could occur (Plasticity Model), the second assumed that only evolutionary change in *M* could occur (Eco-Evolutionary Model), while the third allowed both processes to occur simultaneously (Plasticity + Eco-Evo Model).

The Plasticity Model modifies the existing Supply-Demand model for body size dynamics (DeLong 2012; DeLong & Walsh 2015) and assumes that the rate of change in body size, ^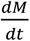^, increases with energy intake (supply), and decreases with energy loss (demand). Following a recent study (DeLong 2020), we assumed that the supply in a species growing logistically depends on the ratio of the carrying capacity, K, and the abundance of the species, N, times a conversion rate constant *e*, that converts the supply to units of *M*. When the population is small, the supply approaches infinity, and it approaches *e* when N grows to K. Following previous studies (DeLong, Hanley & Vasseur 2014), we also assumed that the demand was the metabolic cost of the organism, which is known to increase allometrically with body size, as *cM*^*δ*^. Taken together, the equation controlling changes in body size M was:

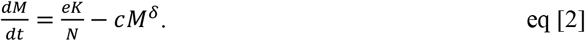

For the Eco-Evolutionary Model, we followed previous studies on eco-evolutionary dynamics (e.g., (Abrams, Harada & Matsuda 1993; Abrams & Matsuda 1997; Jones *et al*. 2009; Ellner & Becks 2010; Jones & Gomulkiewicz 2012; Cortez 2016; Cortez 2018)) to incorporate evolution in *M* at a rate that equals the product of the total amount of additive heritable variation in body size (*i*.*e*., the product of the total phenotypic variance, *σ*^2^, and the narrow sense heritability, h^2^), and the selection gradient (*i*.*e*., the change in fitness, *F*, with respect to a change in *M*, which represents the strength of selection acting on *M*). Taken together, the equation controlling the change in M over time under these assumptions was:

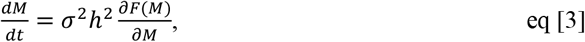

were 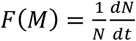 (Lande 1976; Schreiber, Bürger & Bolnick 2011).

Last, the Plasticity + Eco-Evo Model assumes that both Supply-Demand (plastic) and Eco-Evo (rapid evolution) contributions can simultaneously influence the rate of change of *M*, resulting in the following model for body size dynamics:

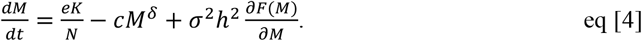

This model does not account for possible interplay between plastic and evolutionary change, such as plasticity facilitating evolution, plasticity impeding evolution, or evolving plasticity, all of which can occur and have been reviewed elsewhere (Diamond & Martin 2016).

None of our models accounts for shifts in age or size structure because here we were are specifically interested in mathematically tracking changes in mean body size, not changes in the entire trait distribution, which also requires model formulations that are not amenable to ODE fitting (Chen, Baños & Buceta 2018; Nieto-Acuña *et al*. 2019).

### Model fitting, parameter uncertainty, and model selection

We fitted the models in Eqs 1-4 to the *T. pyriformis* time series using R package FME v1.3.6.1 (Soetaert & Petzoldt 2010). However, non-linear model fitting tends to get stuck in sub-optimal maxima/minima during residual minimization (or similar procedures) and it is often impossible to simultaneously estimate all model parameters, in which case the model is said to be non-identifiable (Motulsky & Christopoulos 2004; Miao *et al*. 2011). The conversion parameters *a* and *b* of our models –which convert from units of *M*^*α*^ and *M*^*γ*^ into units of *r* and *K*, respectively– were not identifiable (i.e., the fitting procedure could not simultaneously estimate them and all other model parameters without yielding negative or other non-sensical parameter values). Because initial model fits suggested values close to 1 and -1 for the allometric parameter *α* and *γ*, respectively, we estimated *a* and *b* from our data by solving *aM*^*α*^ and *bM*^*γ*^ using the observed intrinsic growth rates for the first two days of growth (r ∼ 3.20 day-1), the observed average *K* ∼ 6400 ind/mL, average *M* (∼10^4^!m^3^) obtained from the FlowCam from day 0 to day 4, and setting *α*=1 and *γ*=−1. Doing so resulted in initial parameter values of ∼10^−4^ for *a* and ∼10^7^ for *b*, which were then optimized during preliminary model fitting (i.e., the iterative process of providing initial parameter guesses and assessing model fit to increase the chance that the fitting procedure succeeds). Because parameters *a* and *b* do not play an important biological role –they convert units of body size (to the power of E or F) into units of *r* or *K*– these parameters should not be expected to change across models, and were thus assumed to be equal for all models and set constant during model parameter and uncertainty estimation of all remaining parameters. Last, we assumed that the scaling parameter of the metabolic cost, *δ*, equaled 1, as has been shown to be the case for protists (DeLong *et al*. 2010), despite it being closer to ¾ for metazoans (Gillooly *et al*. 2001; Brown *et al*. 2004; DeLong *et al*. 2010).

Fitting of ordinary differential equations in package FME relies on the Levenberg-Marquardt algorithm for parameter estimation, and a Metropolis-Hastings MCMC procedure for estimation of parameter uncertainty (Soetaert & Petzoldt 2010). Model comparison was done using Akaike Information Criterion (Burnham & Anderson 2002) as:

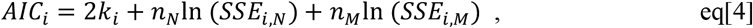

where *k*_*i*_ is the number of parameters of model *i, n*_*N*_ is the number of datapoints considered in the density time series, *n*_*M*_ is the number of datapoints considered in the body size time series, *SSE*_*i,N*_ is the sum of squared errors of model *i* with respect to the density time series, and *SSE*_*i,M*_ is the sum of squared errors of model *i* with respect to the body size time series. An AIC difference >2 indicates that one model is significantly better than the alternative (Burnham & Anderson 2002). Akaike weights were calculated as e^−0.5Δ*AIC*^, where Δ*AIC*=*AIC*_*i*_ − min (*AIC*). Weights are bound between 0 and 1 and models with larger weights can be interpreted as having larger relative likelihoods (Burnham & Anderson 2002). All data and code are available at https://anonymous.4open.science/r/Tetra_Rapid_BodySize_Shifts-5FD7.

## RESULTS

### Time series analyses

Tetrahymena abundances increased to carrying capacity roughly 4 days after microcosms where initialized (Fig 2a, Fig S1a). Body size increased over the first day, then decreased more or less continuously for 12-13 days (Fig 2b, Fig S1b). The CCM analysis showed large predictability values for densities using changes in body size as a predictor, but smaller predictability for body size using densities as the predictor (Fig 2c). The stronger effect of change in body size on density was found to be causal (sharp increase in predictability with library size and decrease in standard deviation, Fig 2c). The effect of density on body size seemed to only be weakly causal (slow convergence of predictability and little change in standard deviation, Fig 2c). This indicated that while changes in body size more strongly influenced density changes, both seemed to have at least some level of influence on each other.

### Experimental results

Our density manipulation resulted in a roughly two-fold difference in density among experimental jars at Day 0 (p<10^−5^, Appendix Fig S2), while the manipulation of body size resulted in a 20% size difference in average body size among experimental jars (p<10^−8,^ Appendix Fig S3). A Tukey post-hoc test indicated no statistically significant differences in body size between high- and low-density jars at Day 0 (p=0.15, Fig 3a), and no statistically significant differences in density between large- and small-size treatments at Day 0 (p=0.64, Fig 3b). Together, these results indicate that we correctly manipulated density and size while keeping the other variable constant. Overall, density increased, and size decreased from Day 0 to Day 2 (Table 1, Fig 3), consistent with trends observed in our time series (Fig 2).

**Table 1:**
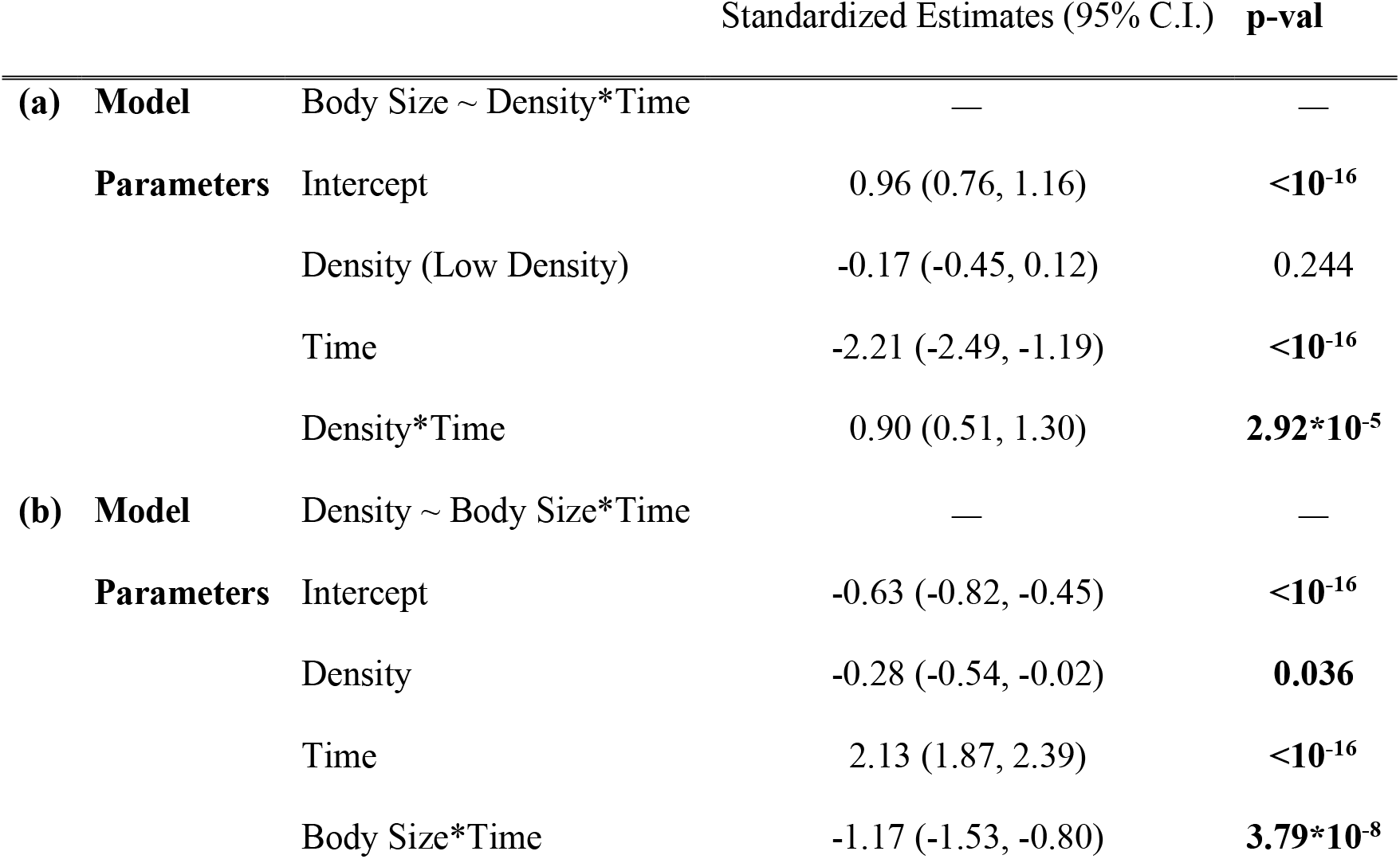
ANOVA results from experimental manipulations of density and size

**Fig 3:**
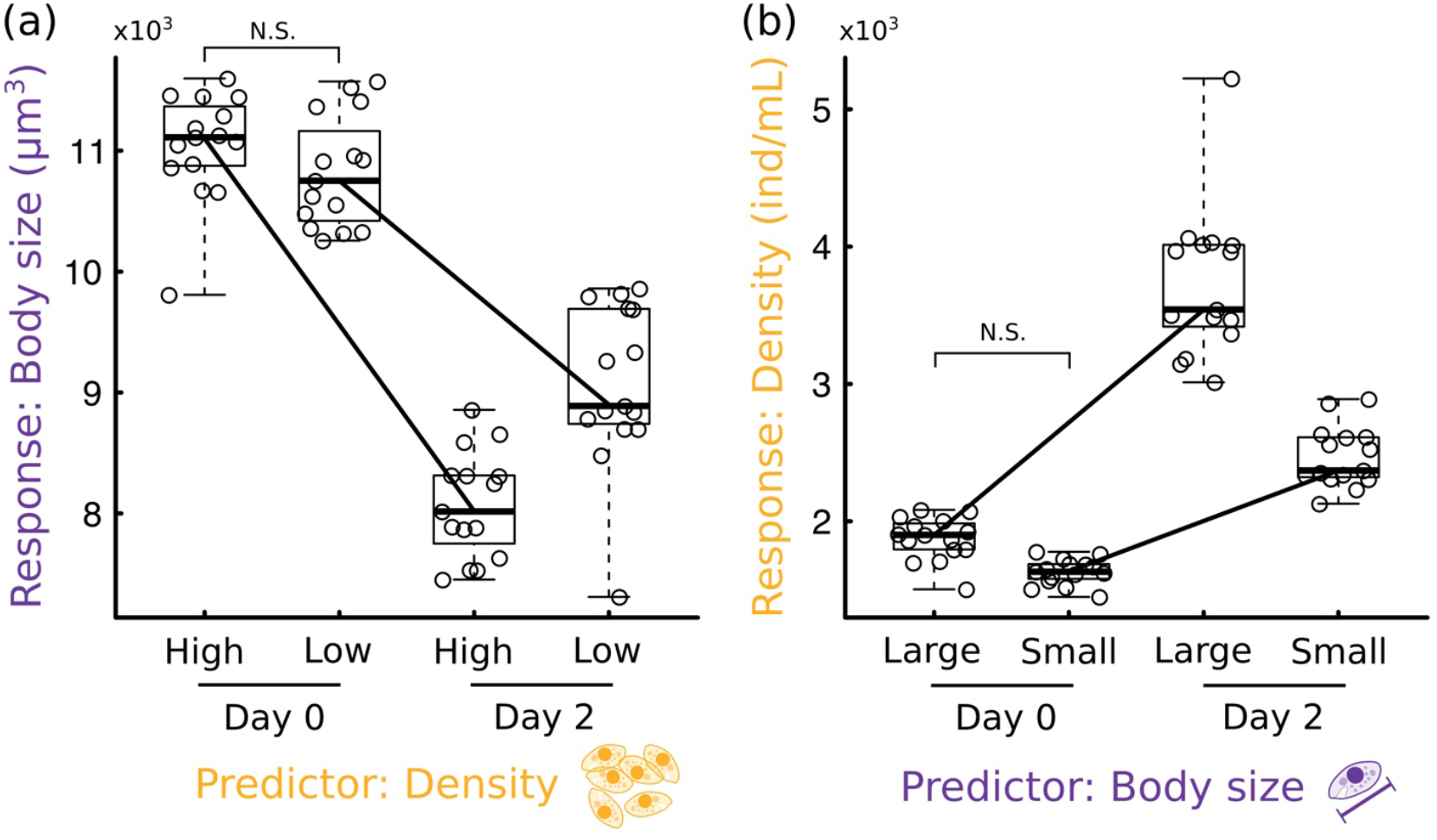
(a) Boxplot of the response of body size over two days across high- and low-density treatments. Solid black lines connecting the bodies of the boxplots are the to help see the interactive effect of the predictor variable and time. (b) Boxplot of the response of density over two days across large- and small-body-size treatments. Solid lines as in (b). N.S. indicates that the manipulation of density (a) and size (b) did not significantly alter the other (size in (a) and density in (b)) at Day 0.

The imposed initial differences in density and size resulted in significant interactions with time (Table 1, Fig 3), indicating significant reciprocal effects of size on density and of density on size. Consistent with results from our CCM analysis (Fig 2c), body size had a larger effect on density (in magnitude) than the other way around (Table 1). However, in standardized units (units of standard deviations, SD), it could be possible for the imposed initial difference in density to be smaller relative to that in size, which could have led to a smaller overall effect of density on size than the other way around. To control for that effect, we divided the observed effect size of density on body size by the imposed (standardized) initial differences in density between low- and high-density jars, and also divided the observed effect size of body size on density by the imposed (standardized) initial difference in body size between small and large size jars. The resulting number could then be interpreted as the magnitude of the effect the predictor variable had on the change observed in the response variable from Day 0 to Day 2 (in units of SD), per unit difference in the initial treatment (also in SD). This resulted in a standardized effect size of body size on density that, while much closer, was still larger in magnitude than that of the effect of density on size (in absolute values, density→size = 0.57, size→density =0.64), still consistent with our CCM results.

### Mathematical models

All models fitted the empirical data remarkably well (Fig 4, Table 2). MCMC chains converged for all fitted parameters (Appendix Fig S4-S9) and model parameters were free of correlations for 25 out of 31 total parameter pairs across the three models (except for *γ* and *m*, and *c* and e in the Plasticity Model, *α* and *m* in the eco-evo model, and c, e and *σ*^2^h^2^, in the Plasticity+Eco-Evo model, Fig S5, S7, and S9). The Plasticity and Plasticity+Eco-Evo Models fit the abundance data slightly better than the Eco-Evo Model (notice departures of Eco-Evo Model in the early time steps, Fig 4a, c, e), while the Plasticity and Eco-Evo Models fit the body size data better than the Plasticity + Eco-Evo Model (Fig 4b, d, f). All models arrived at very similar fitted values for model parameters they had in common (Table 2), indicating good agreement between them all. Consistent with the literature, all models identified the scaling of K and M, *γ*, as a negative number close to -¾ (Table 2). However, all models identified the scaling between r and M, *α*, as positive and close to 1 (Table 2), while the literature pins that value—across species—to -¼ (Savage *et al*. 2004).

**Table 2:**
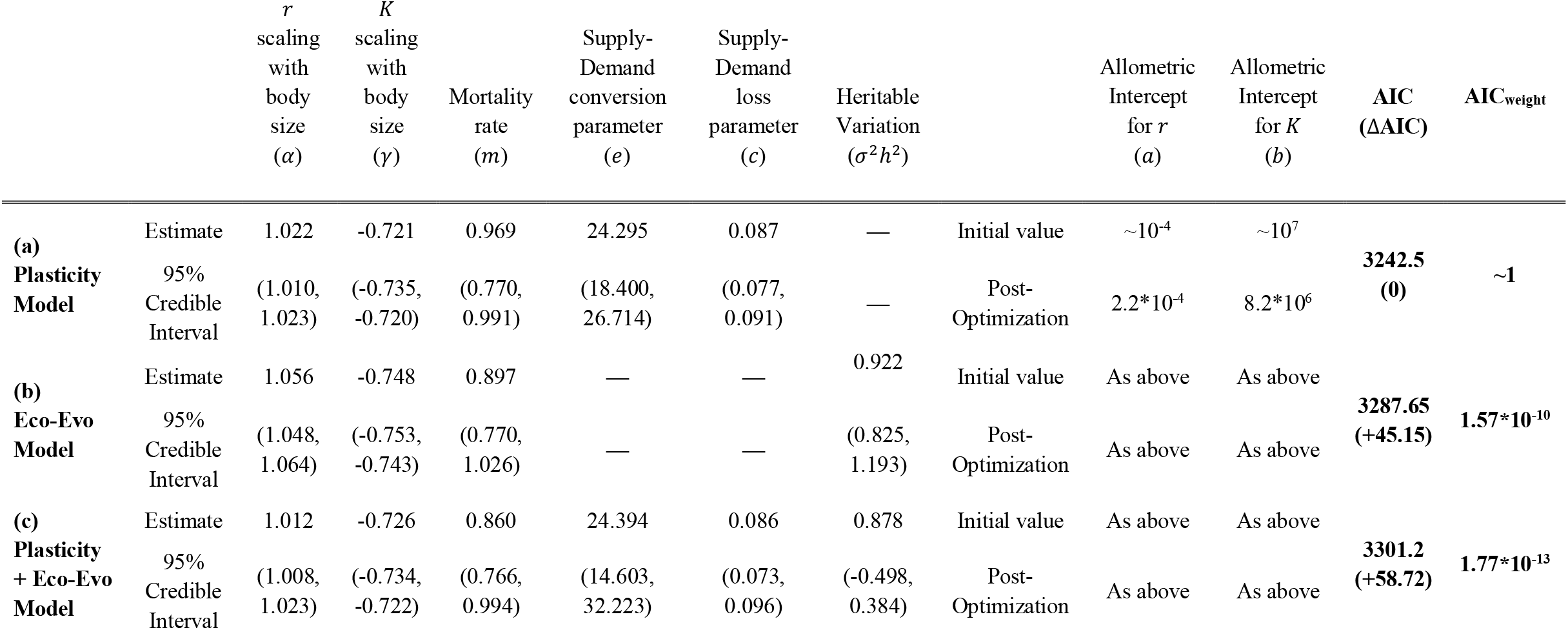
Parameter estimates, parameter uncertainty, and model selection for all fitted models.

| | | *r*<br>scaling<br>with<br>body<br>size<br>($\alpha$) | *K*<br>scaling<br>with<br>body<br>size<br>($\gamma$) | Mortality<br>rate<br>($m$) | Supply-Demand<br>conversion<br>parameter<br>($e$) | Supply-Demand<br>loss<br>parameter<br>($c$) | Heritable<br>Variation<br>($\sigma^2 h^2$) | | Allometric<br>Intercept<br>for *r*<br>($a$) | Allometric<br>Intercept<br>for *K*<br>($b$) | AIC<br>(ΔAIC) | AIC<sub>weight</sub> |
| --- | --- | --- | --- | --- | --- | --- | --- | --- | --- | --- | --- | --- |
| (a)<br>Plasticity<br>Model | Estimate | 1.022 | -0.721 | 0.969 | 24.295 | 0.087 | — | Initial value | $\sim 10^{-4}$ | $\sim 10^7$ | 3242.5<br>(0) | $\sim 1$ |
| | 95%<br>Credible<br>Interval | (1.010,<br>1.023) | (-0.735,<br>-0.720) | (0.770,<br>0.991) | (18.400,<br>26.714) | (0.077,<br>0.091) | — | Post-Optimization | $2.2 \cdot 10^{-4}$ | $8.2 \cdot 10^6$ | | |
| (b)<br>Eco-Evo<br>Model | Estimate | 1.056 | -0.748 | 0.897 | — | — | 0.922 | Initial value | As above | As above | 3287.65<br>(+45.15) | $1.57 \cdot 10^{-10}$ |
|  | 95%<br>Credible<br>Interval | (1.048,<br>1.064) | (-0.753,<br>-0.743) | (0.770,<br>1.026) | — | — | (0.825,<br>1.193) | Post-Optimization | As above | As above |  |  |
| (c)<br>Plasticity<br>+ Eco-Evo<br>Model | Estimate | 1.012 | -0.726 | 0.860 | 24.394 | 0.086 | 0.878 | Initial value | As above | As above | 3301.2<br>(+58.72) | $1.77 \cdot 10^{-13}$ |
|  | 95%<br>Credible<br>Interval | (1.008,<br>1.023) | (-0.734,<br>-0.722) | (0.766,<br>0.994) | (14.603,<br>32.223) | (0.073,<br>0.096) | (-0.498,<br>0.384) | Post-Optimization | As above | As above |  |  |

**Fig 4:**
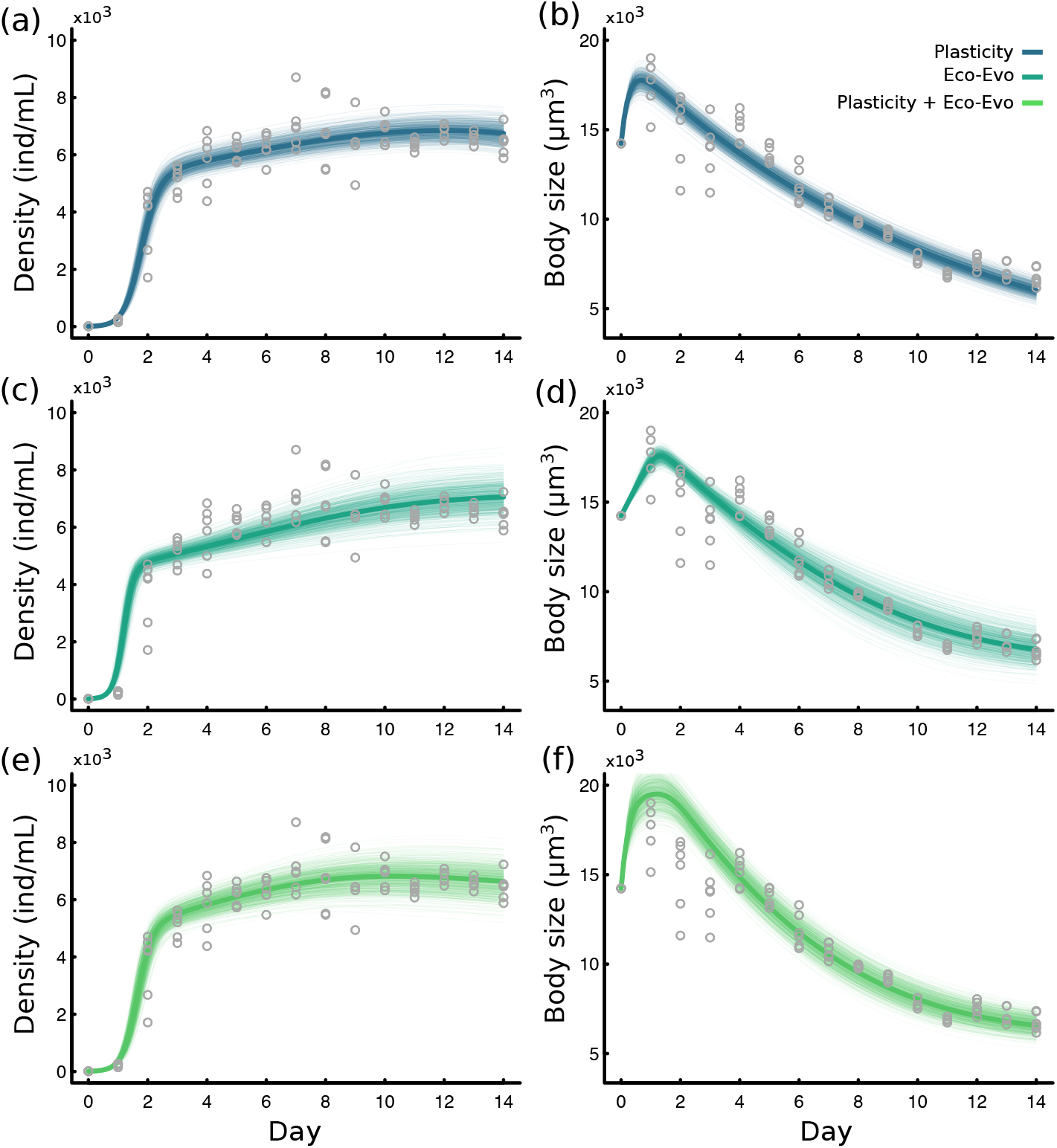
(a) Density data (grey points) and Plasticity Model fit (solid blue). Uncertainty is represented as 700 model predictions (transparent lines) whose parameters were sampled from posterior distributions for each model parameter, estimated during model fitting (Table 2). (b) As in (a), but for body size data. (c–d, e–f) As in (a–b) but for the Eco-Evo Model fit (c–d) or the Plasticity + Eco-Evo Model fit (e–f).

Despite all models fitting the data well, model selection through AIC indicated very large differences in model likelihood, with the Plasticity Model being –by far– the most likely (AIC_weight_ ∼ 1 for the Plasticity Model, but effectively zero for the other two models, Table 2). The Plasticity + Eco-Evo Model was the least likely of all fitted models, perhaps owing to a larger number of model parameters, which are penalized by AIC (Table 2). This result thus suggests that the observed coupled density and body-size dynamics were mostly driven by plasticity, while rapid evolution or a combination of plasticity and rapid evolution are less likely to explain the observed dynamics.

## DISCUSSION

Because of the myriad ecological consequences of body size (e.g. (Gillooly *et al*. 2001; Brown *et al*. 2004; DeLong *et al*. 2010)), it is important to understand how changes in body size—plastic or evolutionary—may influence, or be influenced by, ecological dynamics. We show that changes in body size more strongly influence changes in density than the other way around (Fig 2c, 3b), but that density also influences changes in body size (Fig 2c, 3a). This suggests the existence of a (possibly asymmetric) feedback between the two. Additionally, a model that accounts for rapid plastic change in body size provides the most parsimonious explanation for the observed, coupled ecological and phenotypic dynamics (Fig 4, Table 2). Previous results indicated that phenotypic change often lags ecological change (e.g., (DeLong *et al*. 2016)) but that, under certain conditions, very rapid shifts in body size may precede important changes in ecological dynamics and can thus be used as early warning signals of state shifts (Clements and Ozgul 2015). Our results add to this literature by showing that phenotypic change not only occurs well within ecological timescales and responds to ecological dynamics, but may even causally influence those dynamics (Figs 2, 3).

Understanding the mechanisms of this possible feedback between size and density dynamics is central to gain insights as to how coupled eco-phenotypic dynamics may occur in the wild. In many unicellular organisms, cell growth (increase in body size) at the individual level and cell division are intimately intertwined: cells grow until a critical size is reached, which triggers DNA synthesis and eventual division (Baserga 1968). Larger cells are closer to the critical size threshold that triggers cell division (Jorgensen & Tyers 2004), leading to faster cell division (reproduction) in the next generation, which ultimately results in faster population growth. This link between size and cell division provides a possible explanation for why our results identify changes in size as important drivers of changes in density (Fig 2c, Fig 3). In line with this argument, all models predicted a positive relationship between body size and the intrinsic growth rate, r (*α* ∼ 1, Table 2). This result stands in contrast to empirical data across species and theoretical expectations, which show lower intrinsic growth rates for larger organisms (or *α* < 0, (Savage *et al*. 2004)). Within species, however, larger individuals typically reproduce more and die at lower rates (Peters 1983), leading to higher *r*, due to lower mortality and higher reproduction (Kingsolver & Huey 2008). This positive relationship between body size and the demographic processes that fuel intrinsic growth rates are well understood within species (Peters 1983; Kingsolver & Huey 2008) –emphasizing how inter-species and intra-species body size scaling may often differ (Rall *et al*. 2012)– but also providing a plausible mechanism through which changes in size may be causally influencing changes in density, as our results show.

On the flip side, cells can enter and exit the cell division cycle depending on internal and external cues, such as nutrient availability (Baserga 1968; Fukada *et al*. 2007). As the *T. pyriformis* population reaches carrying capacity, low resource levels likely cue cells to exit the cell division cycle, resulting, in turn, in stunted growth and reduced average body sizes (because cells grow to reproduce) thus providing a possible explanation as to how density may influence body size. If that is the case, then observed total standing phenotypic variation in our population should be largely non-heritable, as also suggested in a recent study (Jacob & Legrand 2021). Surprisingly, the Eco-Evolutionary and Plasticity + Eco-Evo Models support this idea, as they indicate that the total amount of standing heritable variation in body size is rather small (*σ*^2^*h*^2^=0.992 with high confidence for the Eco-Evo model and *σ*^2^*h*^2^=0.878 for Plasticity + Eco-Evo although with very low confidence, Table 2). For comparison, the total amount of phenotypic variation (heritable or not) in our initial population was 94 (units of *μ*m^3^ squared), so the heritable portion of that total phenotypic variation would be on the order of 1%. Shifts in *T. pyriformis* phenotype have also been shown to occur differentially across environmental conditions (DeLong *et al*. 2017; Weber de Melo *et al*. 2020), again suggesting the occurrence of plasticity and very low heritability in this species, and providing support for the above mechanism of response of body size on density.

While the Eco-Evo and Plasticity + Eco-Evo Models were found to be less parsimonious, our results do not rule out the possibility of rapid evolution in this system. Indeed, plastic change often precedes evolutionary change and occurs mostly along the axes of variation with the largest amount of heritable variation (the classic evolutionary ‘path of least resistance’ (Lande 1976; Lande 1979; Lande & Arnold 1983)), thus setting the stage for evolution to occur along those axes (Noble, Radersma & Uller 2019). As we state in our methods, our current models do not account for such a scenario, which could very well be at play here. Indeed, despite low heritability, the less parsimonious Eco-Evo and Plasticity + Eco-Evo models suggest that short term selection imposed by density-dependence may be strong enough to consistently shift body size over time, which in turn influences population dynamics. Our own data indicate that *T. pyriformis* reproduces at a rate of 3.5-4 new individuals per individual per day. This extremely fast population growth may eventually allow for evolutionary change in body size—provided that selection is strong enough, because of low heritability—even if it lags behind plastic change (Chevin, Lande & Mace 2010; Fox *et al*. 2019).

Interestingly, neither water nor nutrients were replenished during our experiment; both were limited and were likely consistently lost from the system through sampling and respiration. This nutrient impoverishment should lead to a strong decline in carrying capacity over time. Such a decline was not, or was only very weakly, observed (Fig 4). However, shifting the focus from abundances to total biomass shows a different picture: biomass increased with abundance in the first few days, but then declined over time (Appendix Fig S10) likely due to density being roughly constant after day 4 but average body size declining consistently after day 2 (Fig 2b, Fig 4, Fig S10). As resources wane from the system, a rapid decline in nutrient concentration may therefore be selecting for smaller body size (Vanni *et al*. 2009), which results in lower total biomass, but also lower competition (through a reduction in metabolic needs associated with smaller size (Brown *et al*. 2004)), ultimately allowing the population to remain at very high densities despite waning resources and increasingly lower biomass. Our data thus suggest interesting ways in which a rapid plastic changes in body size may allow organisms to regulate population growth and density-dependent factors, even as nutrients become increasingly limited.

### Caveats and concluding remarks

Both the CCM and the experimental results agreed that changes in size had a larger effect on changes in density than the other way around (Fig 2c, Fig 3). Yet, they differed on how much stronger this effect of size on density is. This difference may be due to a couple of reasons. First, CCM infers the magnitude of causal effects of one variable on the other throughout the entire time series. It does not inform at what time, exactly, the effect of one variable is larger than the effect of the other variable (Sugihara *et al*. 2012). So, it could very well be that the effect of density is much larger in the first few days –when density changes the most– but then declines over time. Second, our manipulations of size and density likely cannot be extrapolated beyond the first few days of the ecological dynamics (Days 0 to 7), and those first few days coincide with the time span over which larger changes in density were observed. So, while the CCM may underestimate large, temporally localized effects of density (by looking at the entirety of the time series), our experimental work may be overestimating the overall effects of density (as it focuses on their possibly larger effects in the first few days of the dynamics) even though, taken together, both results agree qualitatively.

Because neither CCM nor the experimental work should be used to infer how the effect of size and density on one another may change over time, it is entirely possible, even likely, that both effects change in magnitude over time. This likely explains why, despite body size clearly having an effect on density (Fig 2, 3), density changed little after Day 6 while body size declined from Day 2 to Day 14 (Fig 2b). Indeed, as cells are cued into exiting the cell division cycle due to lack of nutrients, fewer and fewer of them remain reproductive and the proposed mechanism through which cell size may influence density (i.e., through its effect on reproduction) may decline as the population remains at high density and resource scarcity sets in. Taken together, while our results indicate the existence of a feedback between changes in size and changes in density, they also suggest the possibility that the magnitude –and perhaps even the direction of that feedback– may change over time, certainly a promising avenue for future research.

Overall, our study shows that feedbacks between rapid plastic change in body size and change in density are likely integral to the process of population growth itself. This study sheds light on the ecological and evolutionary constraints that regulate population growth and provides new insights about how organisms cope with the negative effects of density-dependence. Our results also emphasize the need to further study and understand the ecological consequences of rapid plastic phenotypic change (Yamamichi, Yoshida & Sasaki 2011; Tariel, Plénet & Luquet 2020), as plasticity, particularly in body size, may play a crucial role in determining the fate of networks of species interactions in a warming world (Barbour & Gibert 2021; Jacob & Legrand 2021).

