## Appendix for "Feedbacks between size and density determine rapid eco-phenotypic dynamics"

INDEX

Fig S1 ………………………………………………………………………………. 1

Fig S2 ………………………………………………………………………………. 2

Fig S3 ………………………………………………………………………………. 3

Fig S4 ………………………………………………………………………………. 4

Fig S5 ………………………………………………………………………………. 5

Fig S6 ………………………………………………………………………………. 6

Fig S7 ………………………………………………………………………………. 7

Fig S8 ………………………………………………………………………………. 8

Fig S9 ………………………………………………………………………………. 9

Fig S10 ………………………………………………………………………………. 10

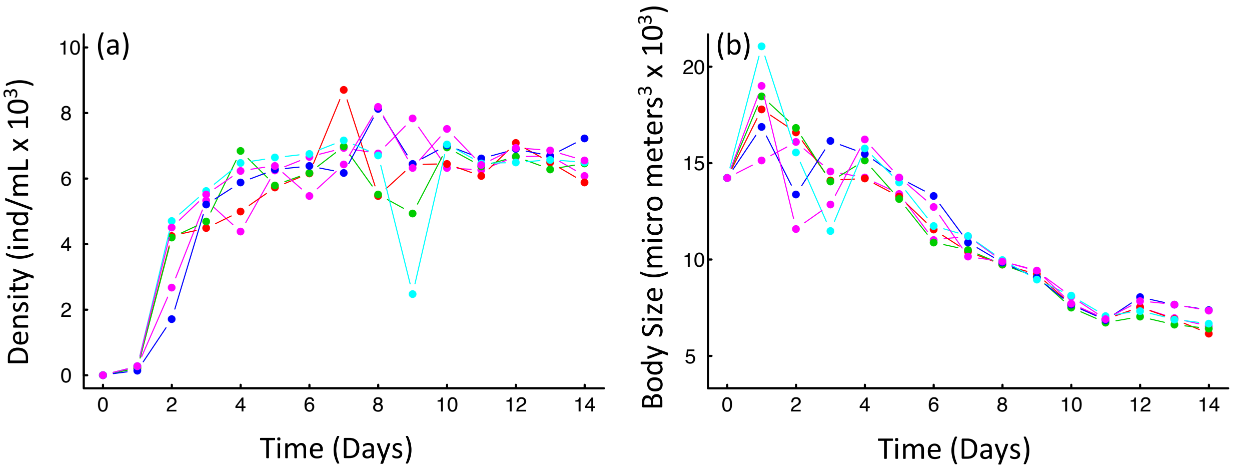


Fig S1: (a) Change in Density over time color coded by jar. (b) Change in Body size over time, color coded by jar with color matching (a) for comparison.

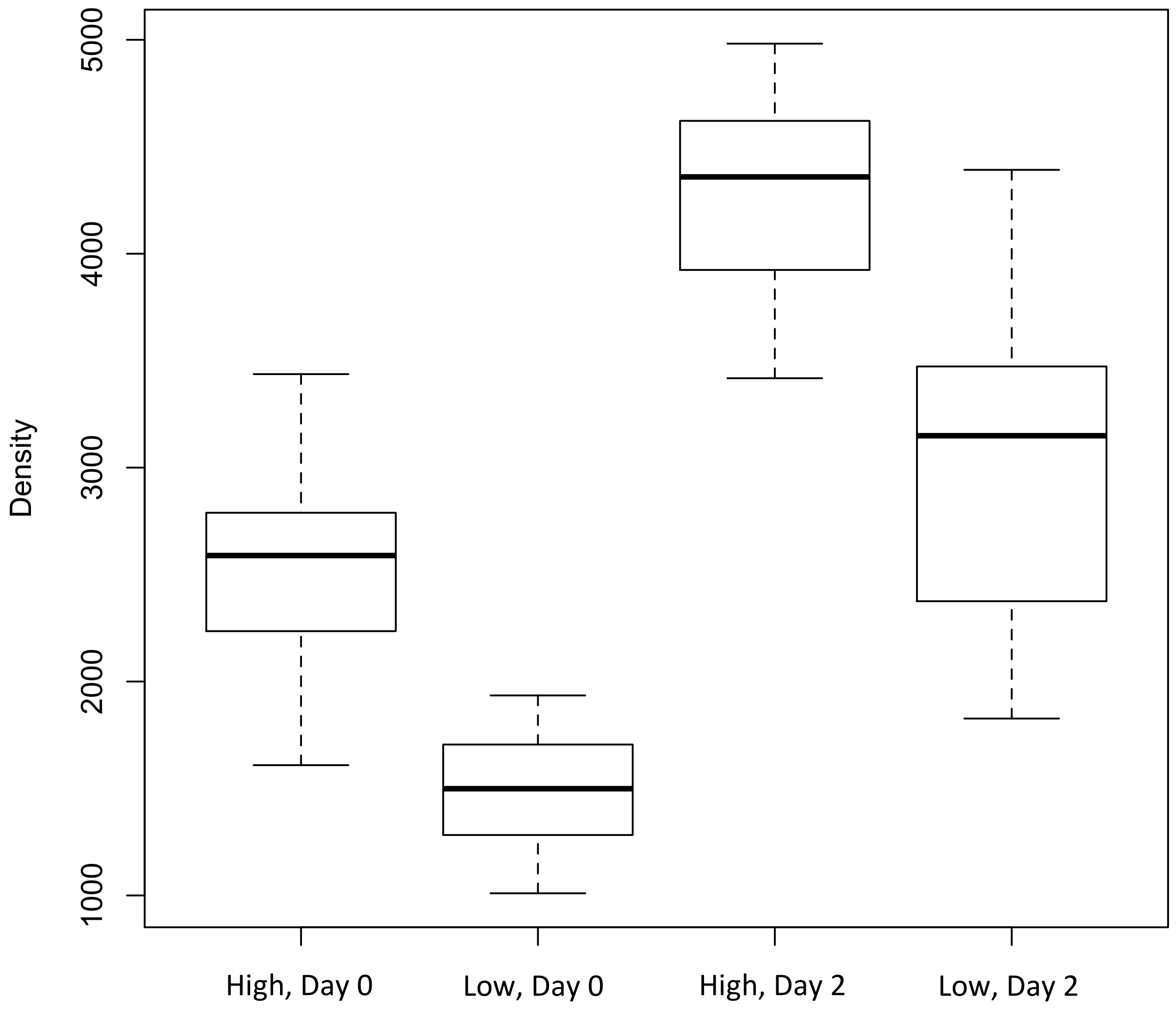


Fig S2: Density difference between high and low density treatments at Day 0 and Day 2. Results from post-hoc Tukey indicate that the difference imposed by the treatment at Day 0 was significant ($p<10^{-5}$). This indicates that our treatment did in fact impose a significant difference in density, but not size (Fig 3 of main text), as intended.

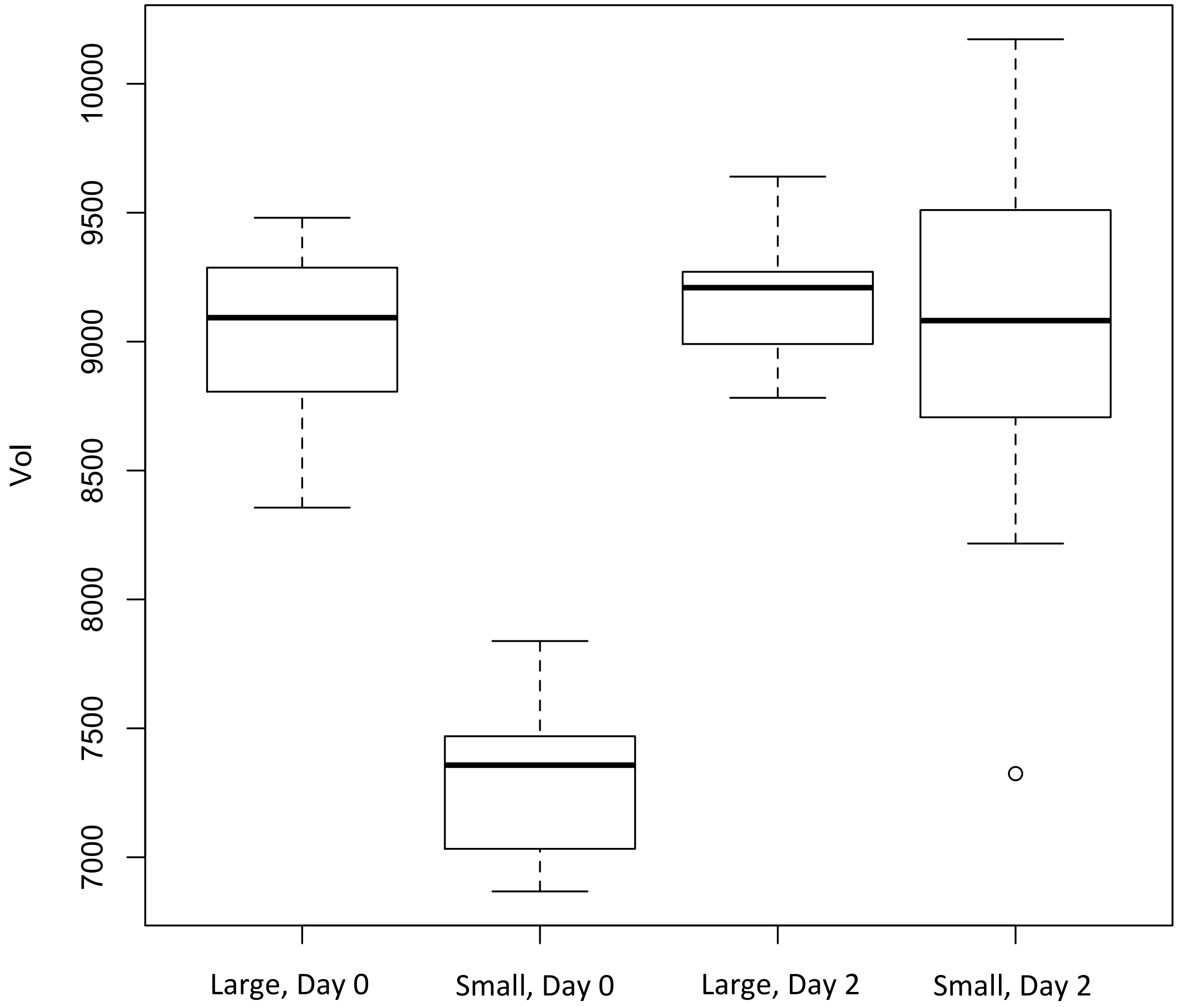


Fig S3: Body size difference between large and small size treatments at Day 0 and Day 2. Results from post-hoc Tukey indicate that the difference imposed by the treatment at Day 0 was significant at ($p<10^{-8}$). This indicates that our treatment did in fact impose a significant difference in size, but not density (Fig 3 of main text), as intended.

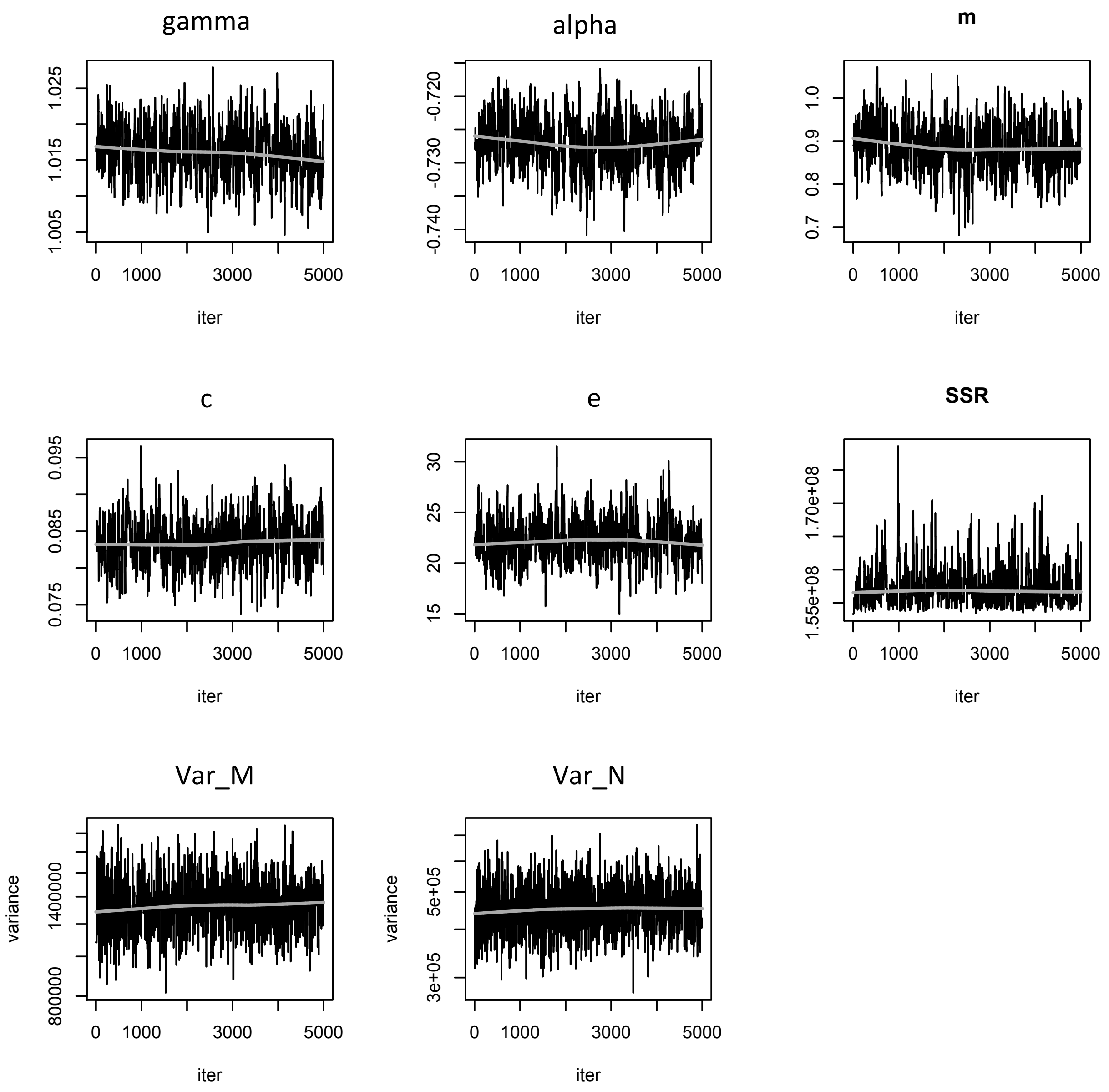


Fig S4: MCMC chains for the plasticity model. Reference: SSR is the sum of squared residuals of each model, Var_M is the variance in body size, and Var_N the variance in density.

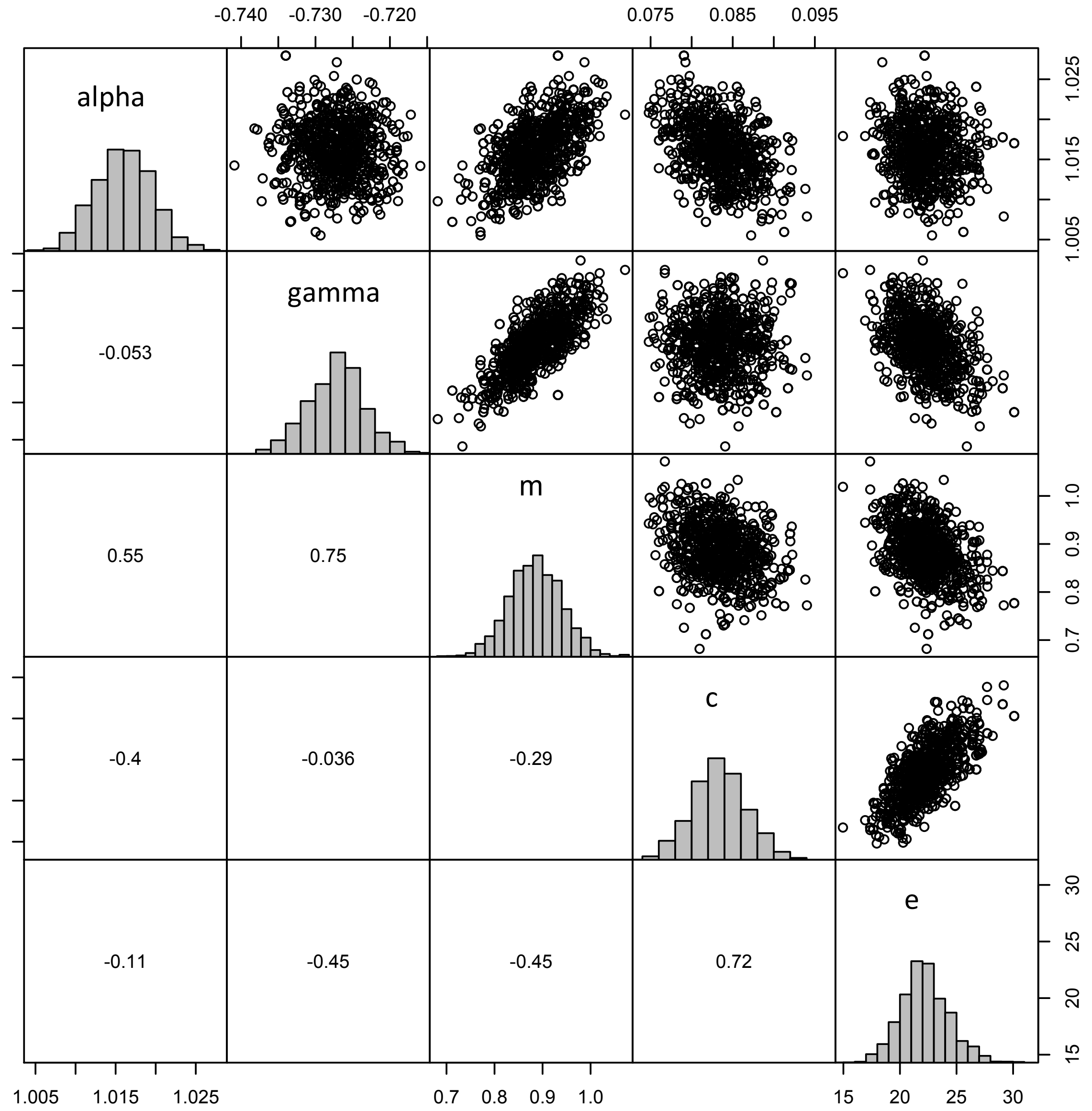


Fig S5: Posterior distributions for all fitted parameters and correlation between parameters.

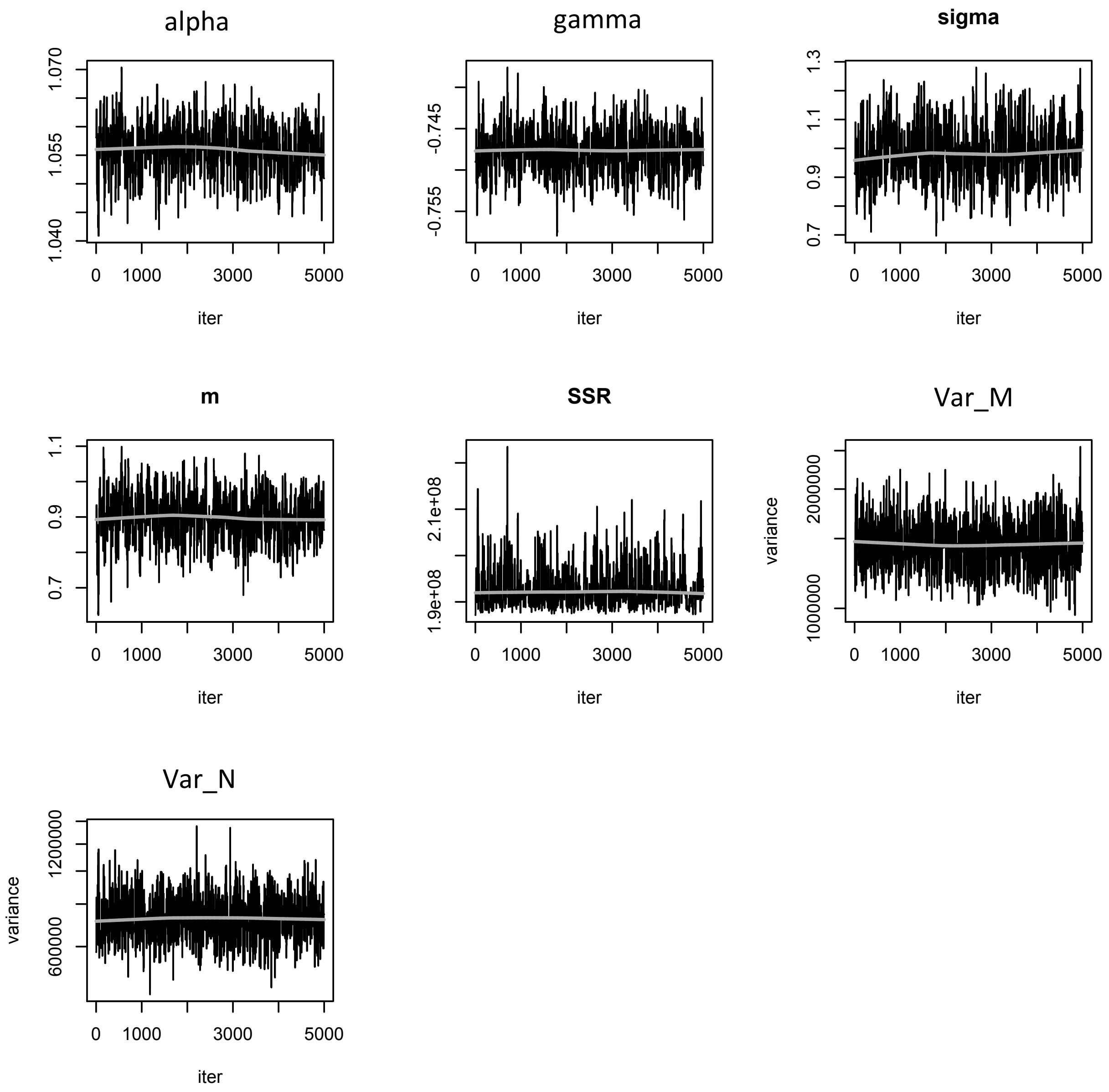


Fig S6: MCMC chains for the eco-evo model. Reference: SSR is the sum of squared residuals of each model, Var_M is the variance in body size, and Var_N the variance in density.

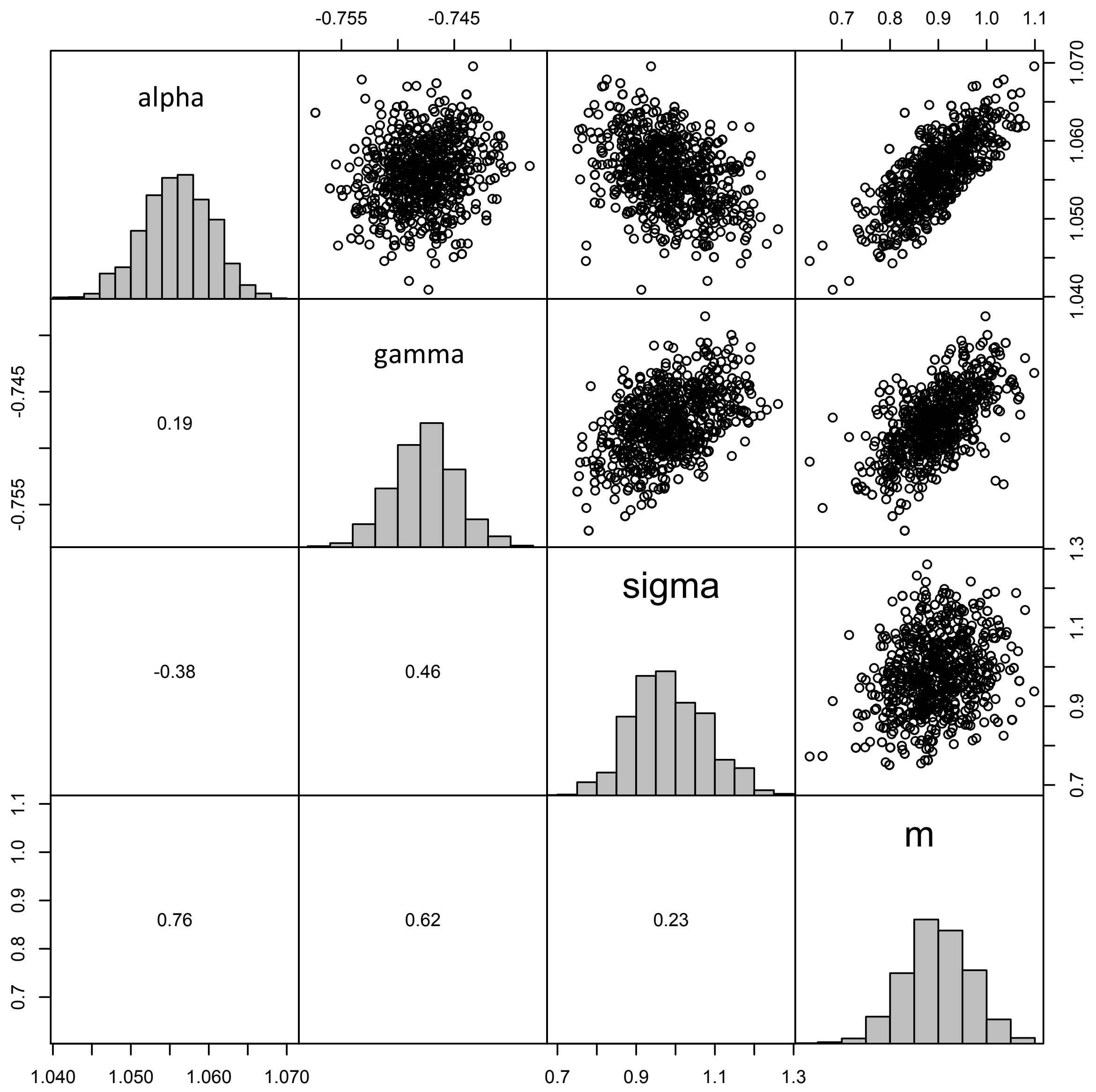


Fig S7: Posterior distributions for all fitted parameters and correlation between parameters.

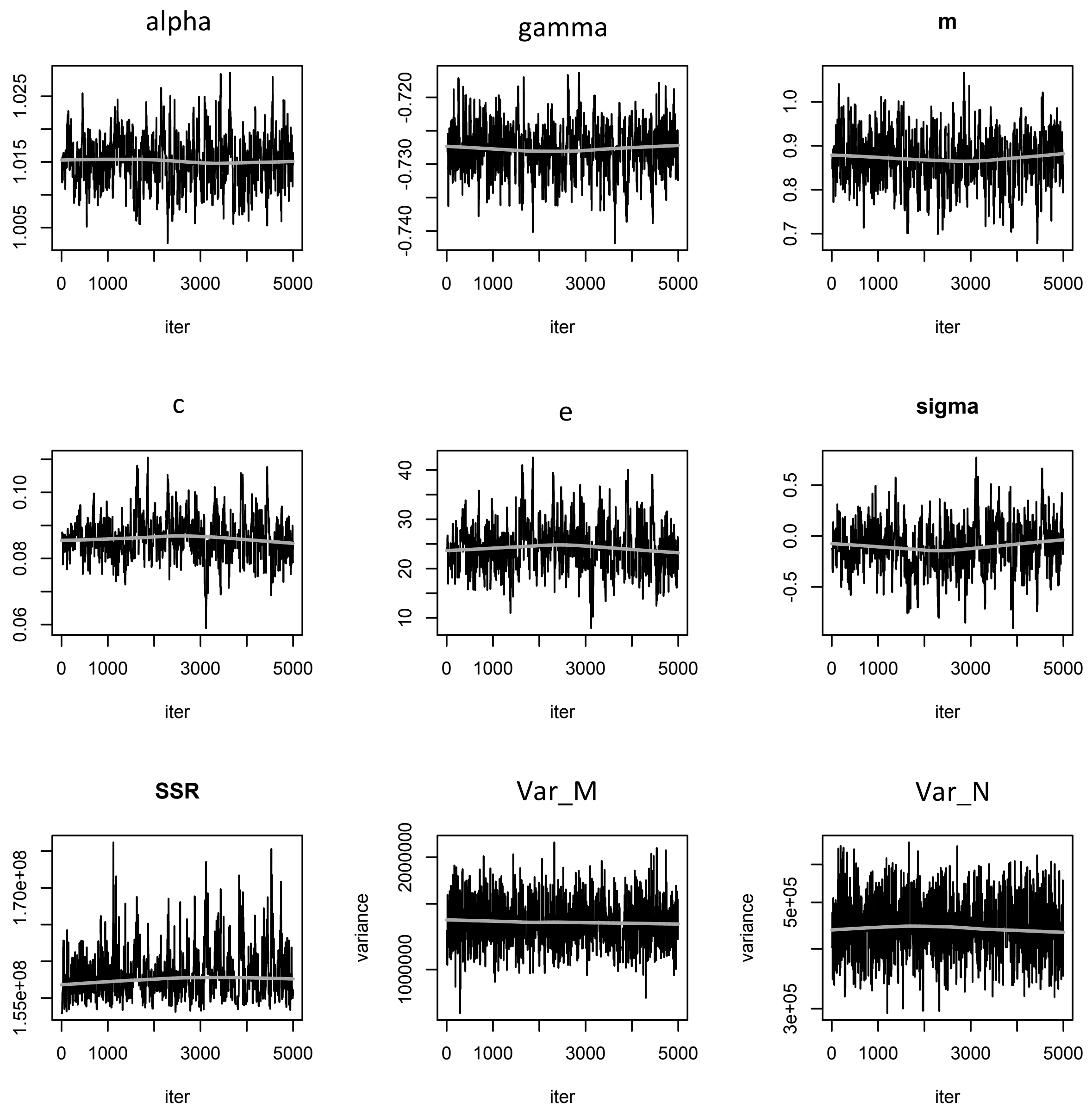


Fig S8: MCMC chains for the eco-evo model. Reference: SSR is the sum of squared residuals of each model, Var_M is the variance in body size, and Var_N the variance in density.

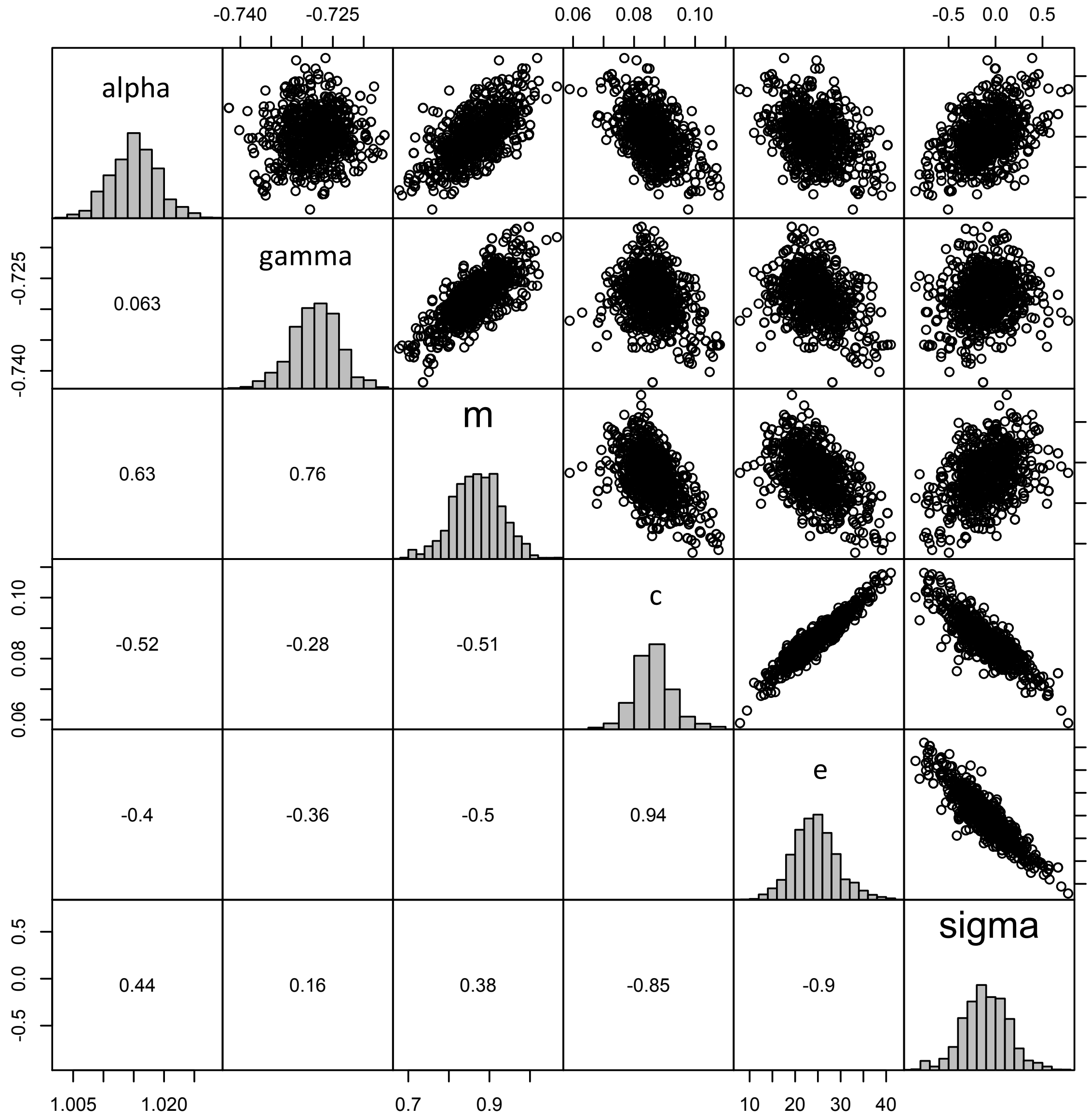


Fig S9: Posterior distributions for all fitted parameters and correlation between parameters. Stronger correlations between parameters c, r and sigma indicate that the model has a harder time providing independent estimates for the three.

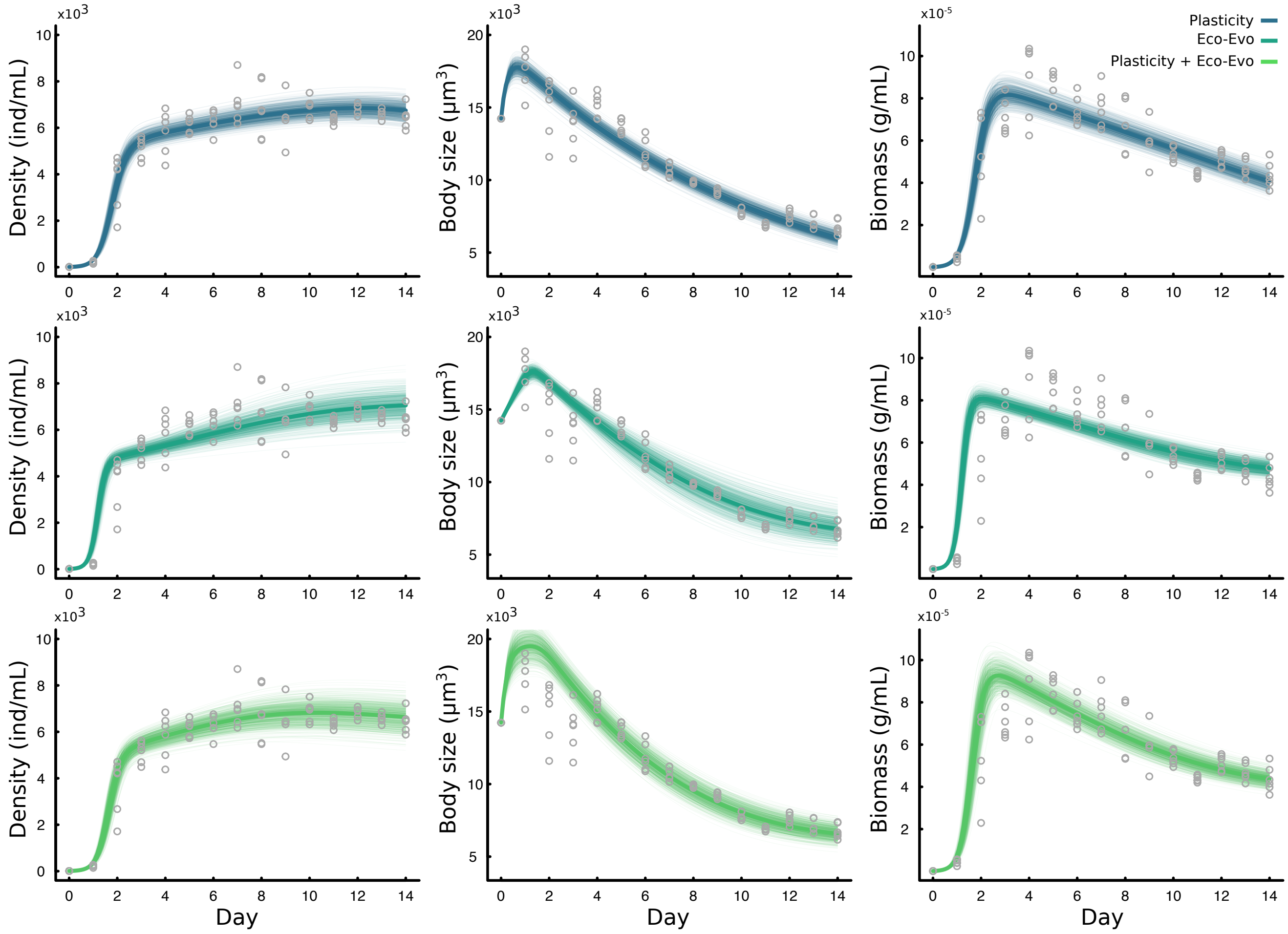


Fig S10: Left and center columns are as in Fig 4 of the main text. The right most column shows how total biomass (in g/mL) changes over time and how each model predicted change in biomass overlays on the actual biomass data (calculated as the product of N(t)*M(t) after fitting the model to the Density and Body Size data).
